# Maternal iron deficiency elicits divergent cardiac mitochondrial responses in normotensive and hypertensive pregnancy

**DOI:** 10.64898/2026.05.12.724698

**Authors:** Jad-Julian R. Rachid, Claudia D. Holody, Si Ning Liu, Rohini Roy Roshmi, Navdeep Badhan, Anson Wong, Alyssa Wiedemeyer, Jennie Vu, Ibrahim Khodabocus, Hélène Lemieux, Stephane L. Bourque

## Abstract

Maternal iron deficiency (ID) during pregnancy is associated with cardiovascular adaptations, including reduced blood pressure and improved cardiac efficiency in hypertensive pregnancy. However, whether these functional gains are accompanied by preserved mitochondrial function remains unclear. Given the metabolic demands of the heart and iron’s central role in oxidative metabolism, we examined how maternal ID affects cardiac mitochondrial ultrastructure, respiration, dynamics, and redox status in pregnant spontaneously hypertensive rats (SHR) and normotensive Wistar-Kyoto (WKY) rats. Female SHR and WKY rats were fed iron-replete or iron-restricted diets before and throughout gestation. On gestational day 21, mitochondrial ultrastructure was assessed by transmission electron microscopy, respiration by high-resolution respirometry, mitochondrial dynamics and quality control proteins by immunoblotting, and antioxidant gene expression by RT-qPCR. Iron restriction reduced maternal haemoglobin levels in both strains. ID dams exhibited enlarged mitochondria with reduced cristae density and lower succinate-supported respiration. SHR dams exhibited reduced fusion signalling, reflected by a lower L-OPA1:S-OPA1 ratio, lower MFN2 abundance, and ID-associated reductions in MFN1 and MFN2. In contrast, DRP1 phosphorylation increased in ID-WKY dams. Iron restriction increased LC3-II:I ratio and BNIP3 in SHR, increased PINK1 in both strains, and increased antioxidant gene expression in ID-SHR but decreased in ID-WKY dams. Despite these alterations, downstream apoptosis activation was not observed. Maternal ID was associated with ultrastructural remodelling and constrained iron-dependent respiration in both strains, with strain-specific alterations in dynamics, mitophagy, and redox signalling, suggesting that favourable haemodynamic adaptations to maternal IDA in hypertensive pregnancy may coexist with underlying bioenergetic constraints in the maternal heart.

**Clinical Perspective:**

- Iron deficiency anaemia is common during pregnancy and may alter how the maternal heart adapts to hypertension, but its effects on cardiac mitochondrial health remain unknown.
- We show that maternal iron deficiency alters cardiac mitochondrial structure and selectively limits iron-dependent respiration in pregnant rats, with different mitochondrial dynamics and stress responses in normotensive and hypertensive strains.
- When considered alongside the previously reported haemodynamic phenotype, these findings indicate that apparently favourable maternal cardiovascular performance can coexist with underlying myocardial bioenergetic constraints, with potential implications for cardiovascular vulnerability during pregnancy and postpartum recovery.

## Introduction

Hypertension is a common complication of pregnancy, affecting approximately 16% of pregnant individuals worldwide each year (1, 2). Hypertensive disorders of pregnancy (HDP), encompassing gestational hypertension, chronic hypertension, preeclampsia/eclampsia, and chronic hypertension with superimposed preeclampsia, are a leading cause of maternal and perinatal morbidity and mortality (2). Anaemia is another highly prevalent gestational complication, affecting approximately 37% of pregnancies globally (3, 4). Iron deficiency (ID), the leading nutritional deficiency worldwide, is the primary cause of anaemia. ID anaemia (IDA) is associated with adverse maternal and fetal outcomes, including maternal haemorrhage, preterm birth, intrauterine growth restriction, and long-term developmental programming of disease risk (5, 6). Despite its clinical prevalence, the cardiovascular consequences of maternal IDA in hypertensive pregnancy remain poorly understood.

We recently demonstrated that mild maternal IDA induces seemingly favourable cardiovascular adaptations in the pregnant spontaneously hypertensive rat (SHR) (7). Specifically, IDA progressively lowered blood pressure throughout gestation while improving cardiac efficiency and ventricular-arterial coupling without impairing uterine blood flow (7). These findings are consistent with the haemodynamic consequences of anaemia, including reduced blood viscosity and systemic vascular resistance, which together promote a hyperdynamic circulatory state characterized by reduced afterload and elevated cardiac output (7, 8). However, unlike other forms of anaemia, IDA also imposes the metabolic consequences of ID itself, superimposing impaired iron availability onto the cardiovascular adaptations to reduced circulating haemoglobin (Hb) concentration.

Iron is essential for mitochondrial bioenergetics and redox homeostasis, serving as a critical cofactor for electron transport system (ETS) complexes and antioxidant enzymes (9, 10). Consequently, ID has been associated with mitochondrial structural remodelling, impaired oxidative phosphorylation, and altered redox balance (11). Hypertension is similarly linked to mitochondrial dysfunction, including disrupted dynamics, oxidative stress, and impaired quality control pathways (12). The combined effects of maternal IDA and hypertension during pregnancy may exacerbate myocardial mitochondrial stress, potentially compromising maternal cardiac metabolic adaptations despite apparent systemic haemodynamic benefits. To address this, we extended our prior haemodynamic characterization of maternal IDA in hypertensive pregnancy by examining the myocardial mitochondrial, redox, and cell survival consequences in pregnant SHR and Wistar Kyoto (WKY) dams. This design allowed us to determine not only whether maternal IDA reshapes mitochondrial biology alongside the previously reported haemodynamic changes, but also whether the pregnant hypertensive heart engages a distinct mitochondrial response to iron restriction compared with its normotensive counterpart.

## Methods

### Animals and Treatments

The experiments described in this study were approved by the University of Alberta Institutional Animal Care and Use Committee (reference animal use protocol #974) and conducted in accordance with Canadian Council on Animal Care guidelines. The haemodynamic and cardiac functional outcomes of this cohort were previously reported (7); all mitochondrial and molecular analysis present here are new.

Female WKY and SHR rats (7-8 weeks old) were purchased from Charles River (Kingston, NY) and housed at the University of Alberta animal care facility under a 12-hour light/dark cycle at 23°C with ad libitum access to food and water. Four weeks prior to breeding, WKY and SHR females were randomly assigned to either an iron-deficient diet containing 3 mg iron/kg (ID-WKY, ID-SHR; D03072501) or an iron-replete control diet containing 37 mg iron/kg (Ctl-WKY, Ctl-SHR; D10012G). All purified diets were based on the AIN-93G formula (Research Diets Inc., New Brunswick, NJ, USA) and differed only in ferric citrate content. After the four-week pre-treatment, females were co-housed with strain- and age-matched males fed standard rodent chow (5L0D; PicoLab, St. Louis, MO) for breeding. Pregnancy was confirmed by the presence of sperm in a vaginal smear the following morning, designated as gestational day (GD) 0. Dams remained on their assigned diet throughout pregnancy.

Maternal body weight, food intake, and haemoglobin (Hb) levels were monitored weekly. Hb was measured using a haemoglobinometer (HemoCue 201+ system) from blood collected via saphenous venipuncture. On GD 21, dams were euthanized by exsanguination and excision of the heart. The atria were removed, and ventricular tissue was weighed and either used fresh for mitochondrial respiration experiments (described below), flash-frozen in liquid nitrogen, or embedded in Tissue-Tek optimal cutting medium and subsequently frozen. All tissues were stored at -80 °C until subsequent analysis.

A total of 52 dams (12 Ctl-WKY, 13 ID-WKY, 14 Ctl-SHR, 13 ID-SHR) were used across the experiments described in this study. Not all endpoints were assessed in every animal; per-experiment sample sizes are indicated in the corresponding figure legends.

### Transmission electron microscopy

Inferolateral mid-sections of the left ventricle (LV) and renal cortical samples were fixed overnight at 4°C in a fixative composed of 2.5% glutaraldehyde and 2% paraformaldehyde in 0.15 M cacodylate buffer. Samples were rinsed with 1X PBS and post fixed for one hour with 2% osmium tetroxide in 0.1M phosphate buffer and washed in double distilled water (ddH2O). The samples were then en bloc stained with 1% uranyl acetate in ddH2O and dehydrated with a series of ethanol (30%, 50%, 70%, 90%, and 100%), followed by overnight infiltration of Spurr resin (EM0300, Millipore Sigma), according to the manufacturer’s instructions. The samples were orientated in a mould and polymerized at 65-70°C for 24 h. Resin blocks were ultrathin sectioned (∼100nm) with a Leica Ultra microtome (Leica, EM UC6) with a diamond knife. Images were subsequently acquired at 30K magnification by a blinded operator using a JEOL transmission electron microscope with a LaB6 filament (Model: 2100F) at 200 kV acceleration voltage with a Gatan Orius side-mounted camera running DigitalMicrograph Suite Software (Version 3.23, Gatan, USA).

Image analysis was performed by a blinded researcher using ImageJ/Fiji (v. 1.54h; National Institutes of Health, Bethesda, MD). Mitochondrial boundaries were manually outlined, and total area, perimeter, and internal electron-lucent space were quantified using consistent threshold settings. Cristae density was calculated as one hundred minus the percentage of electron-lucent space within the mitochondrial boundary. Ten images were analyzed per animal (n=4/group), yielding approximately 30-50 mitochondria per biological replicate. Measurements were averaged by animal before statistical analysis.

### Mitochondrial function

High-resolution respirometry was performed at 37°C using three OROBOROS O2k oxygraph systems (Oroboros Instruments, Innsbruck, Austria), as previously described (6, 13, 14). Respiratory capacity of the electron-transferring flavoprotein-supported pathway (FAO-pathway), NADH-linked pathway (N-pathway), succinate-linked pathway (S-pathway), and combined NADH- and succinate-linked pathway (NS-pathway) was assessed using two substrate– uncoupler–inhibitor titration (SUIT) protocols. The FAO-pathway protocol was performed using permeabilized cardiac fibres and renal cortical homogenates, whereas N-, S-, and NS-pathway protocol was performed using cardiac and renal cortical homogenates. The transition from permeabilized cardiac fibres to cardiac homogenate in the second SUIT protocol was implemented to improve assay consistency and reproducibility. All reagents were purchased from MilliporeSigma unless otherwise stated.

For permeabilized cardiac fibres, fresh ventricular tissue on GD21 was immediately placed in ice-cold biopsy preservation solution (BIOPS) (13). Cardiac fibres were mechanically permeabilized by careful dissection in ice-cold BIOPS using fine forceps, followed by gentle agitation for 30 min on ice in BIOPS supplemented with 0.5 mg/mL saponin. Fibres were subsequently rinsed in mitochondrial respiration medium (MiR05; 110 mM sucrose, 0.5 mM EGTA, 3 mM MgCl_2_, 60 mM K-lactobionate, 20 mM taurine, 10 mM KH_2_PO_4_, 20 mM 4-(2-Hydroxyethyl)piperazine-1-ethanesulfonic acid, 1 g/L fatty acid-free bovine serum albumin, pH 7.1) for 10 min on ice under continuous stirring. Fibres were blotted, weighed, and ∼2–5 mg were added to each O2k chamber containing MiR05 at a final volume of 2 mL.

Fresh cardiac ventricular and renal cortical tissue used for homogenate preparation were immediately placed in ice-cold MiR05. Approximately 20–30 mg of tissue was rinsed, blotted, weighed, minced using cold scissors, and homogenized in MiR05 using a Potter–Elvehjem tissue homogenizer at low speed for 2–3 passes. A total of 100 µL of homogenate were added to O2k chambers containing 1.9 mL MiR05.

For the FAO protocol, LEAK respiration was measured in the presence of 0.1mM L-malate and 0.2mM octanoylcarnitine (Tocris R&D Systems, 0605). Oxidative phosphorylation (OXPHOS) capacity through the FAO-pathway was subsequently assessed following addition of ADP (2.5 mmol/L) and cytochrome c (10 μmol/L). Sequential addition of malate (2mmol/L), pyruvate (5 mmol/L), glutamate (10 mmol/L), and succinate (10 mmol/L) were then used to evaluate convergent electron flow through FAO-, N- and S-linked pathways. Maximal ETS capacity was assessed by stepwise titration of 10 mM 2,4-dinitrophenol (1 µL increments up to 5 µL), and residual oxygen consumption (ROX) was measured following inhibition of complex III with antimycin A (5 μmol/L). Flux control ratios (FCRs) were calculated by normalizing oxygen flux at each respiratory state to the maximal uncoupled ETS capacity achieved during DNP titration.

For the N-, S-, and NS-pathway protocol, LEAK respiration was measured in the presence of pyruvate (5 mmol/L), malate (2 mmol/L), and glutamate (10 mmol/L). N-pathway OXPHOS capacity (through complex I) was assessed following the addition of ADP (2.5 mmol/L) and cytochrome c (10 μmol/L). NS-pathway OXPHOS capacity (through complexes I and II) was subsequently measured following the addition of succinate (10 mmol/L), whereas isolated S-pathway capacity was determined following inhibition of complex I with rotenone (1 μmol/L). ROX was assessed following inhibition of complex III with antimycin A (5 μmol/L). Complex IV activity was measured using ascorbate (2 mmol/L) and tetramethylphenylenediamine (0.5 mmol/L), followed by sodium azide (25 mmol/L) addition for chemical background correction. FCRs were normalized to maximal ETS capacity supported by convergent electron flow through the NADH- and succinate-linked pathways.

Mitochondrial respiration data were analyzed using the Datlab software (OROBOROS Instruments, Innsbruck, Austria). Following all respirometry experiments, samples were further homogenized using a rotor-stator homogenizer, aliquoted, and stored at -80°C for analysis of citrate synthase (CS) activity assays.

### Citrate synthase activity

CS activity was assessed in cardiac and renal samples as previously described (6, 13), by monitoring the reduction of 5,5’-dithiobis-(2-nitrobenzoic acid) at 412nm using a UV/Vis spectrophotometer (Ultrospec 2100 Pro; Biochrom, Cambridge, MA, USA) maintained at 37°C.

### Gene Expression

Cardiac gene expression was analyzed by quantitative reverse-transcriptase PCR (qRT-PCR) as previously described (7). Total RNA was extracted from frozen ventricular tissues using TRIzol reagent (Life Technologies, Carlsbad, CA) following the manufacturer’s instructions. RNA purity and concentration were assessed by measuring A260/A280 absorbance using a NanoDrop 800 spectrophotometer (Thermo Fisher Scientific). Purified RNA was reverse transcribed into complementary DNA using 5× All-In-One RT Master Mix (Applied Biological Materials Inc., G592) according to the manufacturer’s instructions.

qRT-PCR was performed using a 384-well optical reaction plate with PowerUp SYBR Green Master Mix (Applied Biosystems, A25742) on a LightCycler 480 system (Roche Life Science). Primer sequences (**Table S1**) were purchased from Integrated DNA Technologies (IDT, Coralville, IA). Melt curve analysis was conducted to confirm primer specificity. Relative gene expression was calculated using -ΔΔCt method, with *Actb* expression as the reference gene (7). The stability of *Actb* as a reference gene was verified in cardiac tissue: mean *Actb* Ct values were comparable across all four experimental groups (range: 20.5-21.4), with a coefficient of variation below 4% within each group, consistent with stable expression across strain and iron status.

### Western Blotting

Frozen LV tissue was homogenized in RIPA lysis buffer (Thermo Fisher Scientific, 89900) with Protease/Phosphatase Inhibitor Cocktail (100X) (Cell Signalling Technology, 5872) at a final 1X concentration, using 100mg of tissue per 1mL of buffer. Total protein concentration of lysate was determined by bicinchoninic acid assay (Thermo Fisher Scientific, 23225). A total of 50 µg of protein per well was separated using 8-16% precast polyacrylamide gels (Bio-Rad, 4568105) before being wet transferred onto 0.45 µm PVDF membranes (MilliporeSigma, IPVH00010) at 4°C for 1 hour at 100V. Membranes were blocked in 5% bovine serum albumin (Sigma Aldrich, A9647) or non-fat dry milk (Bio Rad, 1706404), as per manufacturer recommendations, in Tris-buffered saline with 0.05% Tween-20 (TBST) for 2 hours at room temperature.

Membranes were then incubated overnight at 4°C with primary antibodies: AKT (Cell Signalling Technology, 9272S; 1:1000), p-AKT Ser473 (Cell Signalling Technology, 9271S; 1:1000), BNIP3 (Cell Signalling Technology, 3769S; 1:1000), Caspase 3 (Cell Signalling Technology, 9662S; 1:1000), Caspase 7 (Cell Signalling Technology, 9492S; 1:1000), Drp1 (Cell Signalling Technology, 8570S; 1:1000), p-Drp1 Ser616 (Cell Signalling Technology, 4494S; 1:1000), GSK-3β (Cell Signalling Technology, 9315; 1:1000), p-GSK-3β Ser9 (Cell Signalling Technology, 9323; 1:1000), LC3B (Proteintech, 14600-1-AP; 1:1000), Malondialdehyde (MDA; Abcam, ab27642; 1:1000), MFN1 (Proteintech, 13798-1-AP; 1:1000), MFN2 (Proteintech, 12186-1-AP; 1:1000), NRF2 (Proteintech, 16396-1-AP; 1:1000), OPA1 (Proteintech, 27733-1-AP; 1:3500), Parkin (Proteintech, 14060-1-AP; 1:1000), PINK1 (Proteintech, 23274-1-AP; 1:1000), p62 (Abcam, ab56416; 1:1000).

Following primary antibody incubation, membranes were washed three times with TBST and incubated for 1 hour at room temperature with an HRP-conjugated anti-rabbit (Cell Signalling Technology, 7074S; 1:2000) or anti-mouse (Cell Signalling Technology, 7076S; 1:2000) secondary antibodies. Signal detection was performed using an enhanced chemiluminescence substrate (Bio-Rad, 1705061), and images were acquired using a ChemiDoc imaging system (Bio-Rad, 12003153). Band intensity was quantified using Image Studio software v6.0 (Li-Cor Biosciences, Lincoln, NE), and protein expression was normalized either to total protein using a Pierce reversible protein stain (Thermo Fisher Scientific, 24585) or to non-phosphorylated versions of the respective native protein.

### Dihydroethidium fluorescence

Myocardial cytosolic superoxide was assessed in LV cryosections by dihydroethidium (DHE) fluorescence, as previously described (6, 14). LV tissue embedded in Tissue-Tek optimal cutting temperature compound was sectioned at 5 μm using a cryostat at -20°C and mounted onto glass slides. Sections were washed three times for 2 min each in Hank’s Balanced Salt Solution (HBSS), incubated at 37°C for 10 min in HBSS, and then incubated with DHE (30 μmol/L; MilliporeSigma, 37291) for 30 min at 37°C in the dark. Following incubation, sections were washed three additional times in HBSS under light-protected conditions and mounted with coverslips using HBSS. Fluorescence was visualized using a Leica TCS SP5 confocal microscope (excitation 500 nm, emission 582 nm) at 20× magnification by an operator blinded to treatment group. Fluorescence intensity was quantified by a blinded researcher using ImageJ/Fiji (v. 1.54h; National Institutes of Health, Bethesda, MD, USA).

### Caspase Activity

Caspase-9 and Caspase-3/7 activities were measured using commercially available kits (Abcam Inc.; ab65607 and ab39383; respectively) according to the manufacturer’s instructions. Briefly, cardiac LV tissue (100mg) was homogenized in provided lysis buffer and centrifuged at 12,000xg for 20 min at 4°C. Supernatant protein concentration was determined using a bicinchoninic acid assay (Thermo Fisher Scientific, 23225). For each sample, 25 µg of protein (0.5 µg/µL) was loaded in duplicate with 50 µL of 2X Reaction Buffer containing 10 mM dithiothreitol (DTT). Lastly, 5µL of fluorogenic probe (final concentration: 50µM) was added and plates were incubated at 37°C for 1hr. Fluorescence intensity was measured using a plate reader (excitation 400nm; emission 505nm; BioTek Synergy MX).

### Statistical analysis

Results are shown using box-and-whisker plots, with box edges represent the 25th-75th percentiles, central line indicates the median, and whiskers indicating maximum and minimum values. Sample size (n) reflects the number of individual dams or litters assessed. Continuous data were analyzed by two-way ANOVA followed by Holm-Šídák’s post-hoc test. Discrete data (e.g. litter sizes) were assessed by Kruskal-Wallis test with Dunn’s post hoc test. Normality was examined using the Shapiro-Wilk and Kolmogorov-Smirnov tests, and heteroscedasticity was evaluated using Spearman’s test. When assumptions of normal distribution or equal variance were not met, appropriate data transformations or statistical tests were applied. Outliers were detected and excluded using Grubbs’ test. All statistical analyses were performed using Prism 10 (GraphPad Software Inc., San Diego, CA).

## Results

### Maternal and Litter Outcomes

The effects of maternal dietary iron restriction on pregnancy and fetal outcomes in WKY and SHR dams have been previously described (7). Outcomes for the subset of dams and offspring included in the present study are shown in **Fig. S1**. Maternal iron restriction induced a progressive decline in maternal Hb concentrations throughout gestation in both strains (**Fig. S1A, left**), culminating in anaemia by GD21 (**Fig. S1A, right**). In the same cohort, iron-restricted dams have previously been shown to exhibit increased hepatic *Tfrc* and reduced *Hamp1* expression (7), confirming the observed anaemia arises from functional ID rather than other causes. Maternal ID reduced weight gain by GD21, independent of strain (**Fig. S1B**), despite SHR dams exhibiting smaller litter sizes than WKY dams (**Fig. S1C**). Fetal Hb concentrations were lower in ID pups, although an interaction effect indicated lower Hb levels in control (Ctl) SHR fetuses compared with Ctl-WKY fetuses (**Fig. S1D**). ID offspring exhibited fetal growth restriction relative to controls, although SHR fetuses tended to be slightly heavier than WKY fetuses overall.

### Mitochondrial Morphology and Content

TEM of cardiac sections revealed strain- and ID-dependent alterations in mitochondrial ultrastructure (**Fig. 1A**). Mitochondrial size metrics, including area and perimeter, were greater in SHR compared with WKY dams and were further increased by ID in both strains (**Fig. 1B, C**). In parallel, cristae density, quantified as the percentage of mitochondrial area occupied by electron-dense structures, was lower in SHR than in WKY dams and was further reduced by ID, particularly in WKY dams (**Fig. 1D**). CS activity was unaffected by either strain or iron status (**Fig. 1E**), indicating preserved overall citric acid cycle activity despite marked alteration in mitochondrial morphology. Cardiac expression of *Ppargc1a* and *Ppara* was reduced in ID dams of both strains (**Fig. 1F**).

**Figure 1.**
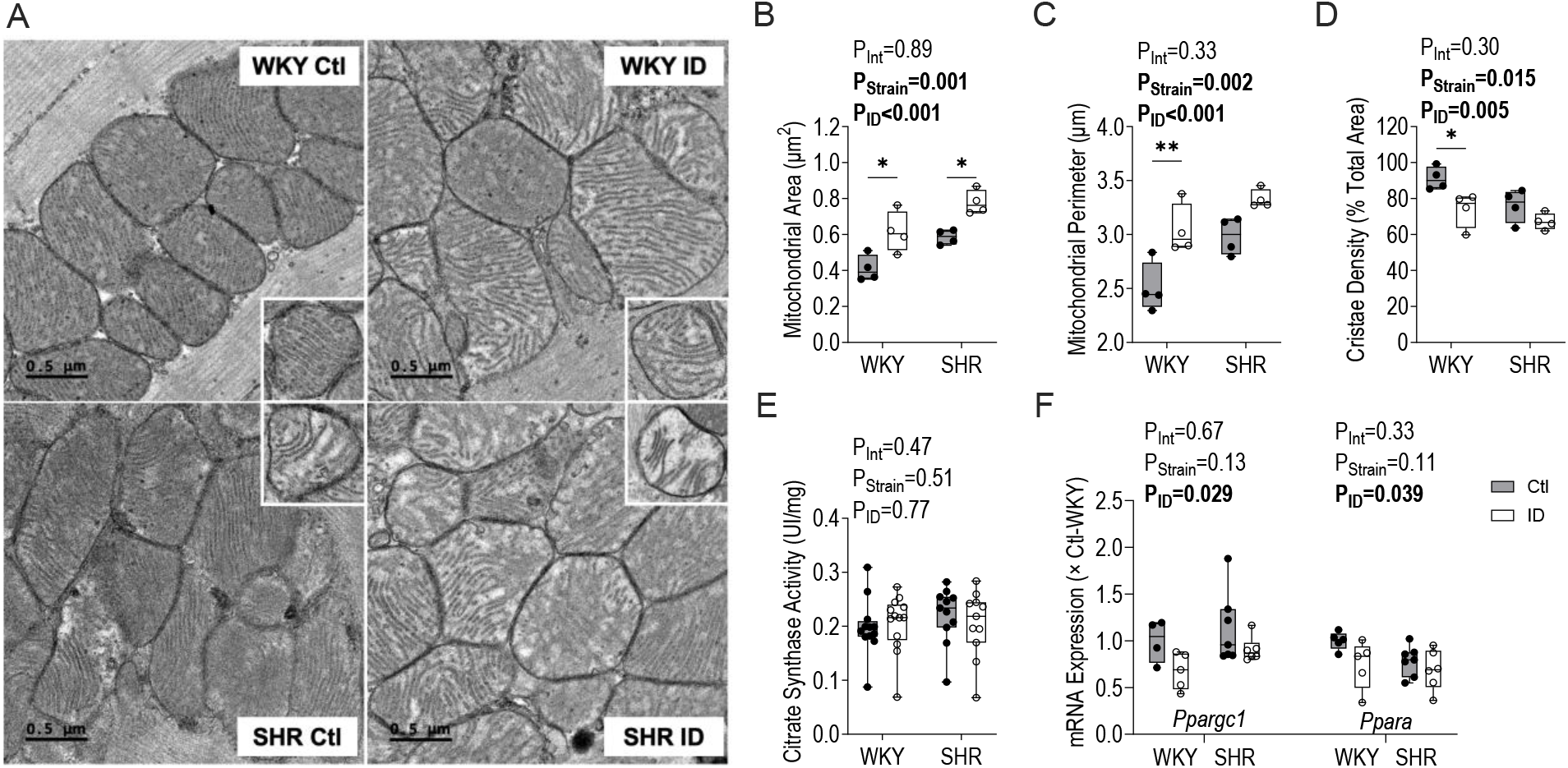
Iron deficiency (ID) alters cardiac mitochondrial ultrastructure in Wistar Kyoto (WKY) and Spontaneously Hypertensive Rat (SHR) dams at gestational day 21. (A) Representative transmission electron microscopy images of left ventricular myocardium (scale bar: 0.5 μm). Inserts show typical ultrastructural features. Quantification of (B) mitochondrial area, (C) mitochondrial perimeter, and (D) cristae density (% total area). (E) Citrate synthase activity (U/mg) in cardiac tissue. (F) mRNA expression of *Ppargc1* (left) and *Ppara* (right), normalized to control (Ctl) WKY. Data are shown as box-and-whisker plots depicting median (middle line), 25th and 75th percentiles (lower and upper edges) and minimum and maximum values (whiskers), with biological replicates overlaid (n=4-13/group in panels B-F). Comparisons were performed using 2-Way ANOVA followed by Holm-Šídák’s post hoc test; *P<0.05, **P<0.01. Statistical outliers, identified using Grubb’s test, included one Ctl-WKY datapoint in panel F (*Ppargc1a*).

To determine whether the observed cardiac mitochondrial phenotype reflected a tissue-specific response to maternal ID, we performed targeted assessments in renal cortical tissue from the same animals (**Fig. S2**). In contrast to the myocardium, cortical mitochondrial area and cristae density were unaffected by strain or ID, although mitochondrial perimeter was greater in SHR compared with WKY dams (**Fig. S2A-D**). Cortical CS activity was similarly unchanged (**Fig. S2E**).

### Mitochondrial Function

Mitochondrial respiratory capacity was assessed in cardiac homogenates by high-resolution respirometry, with oxygen flux normalized to CS activity (**Fig. 2A-D**); Corresponding FCR (i.e. normalized to maximal flux) are presented in **Fig. S3**. N-pathway (through complex I) respiration tended to be lower in ID dams, although this did not reach statistical significance (P_ID_ =0.064; **Fig. 2A**). S-pathway (complex II-linked) respiration was reduced in ID dams, with an effect most apparent in WKY dams (**Fig. 2B**). In contrast, FAO-supported respiration measured in the presence of octanoylcarnitine was unchanged by either strain or iron status (**Fig. 2C**). Complex IV activity was similarly unaffected by strain or ID (**Fig 2D**). Flux control ratios (FCR; **Fig. S3**) were unchanged across the N-, S-, FAO-, and CIV pathways, indicating that despite reductions in absolute respiratory capacity normalized to CS activity, the relative contribution of S-pathway to total respiratory flux remained preserved (**Fig. S3A-D**).

**Figure 2.**
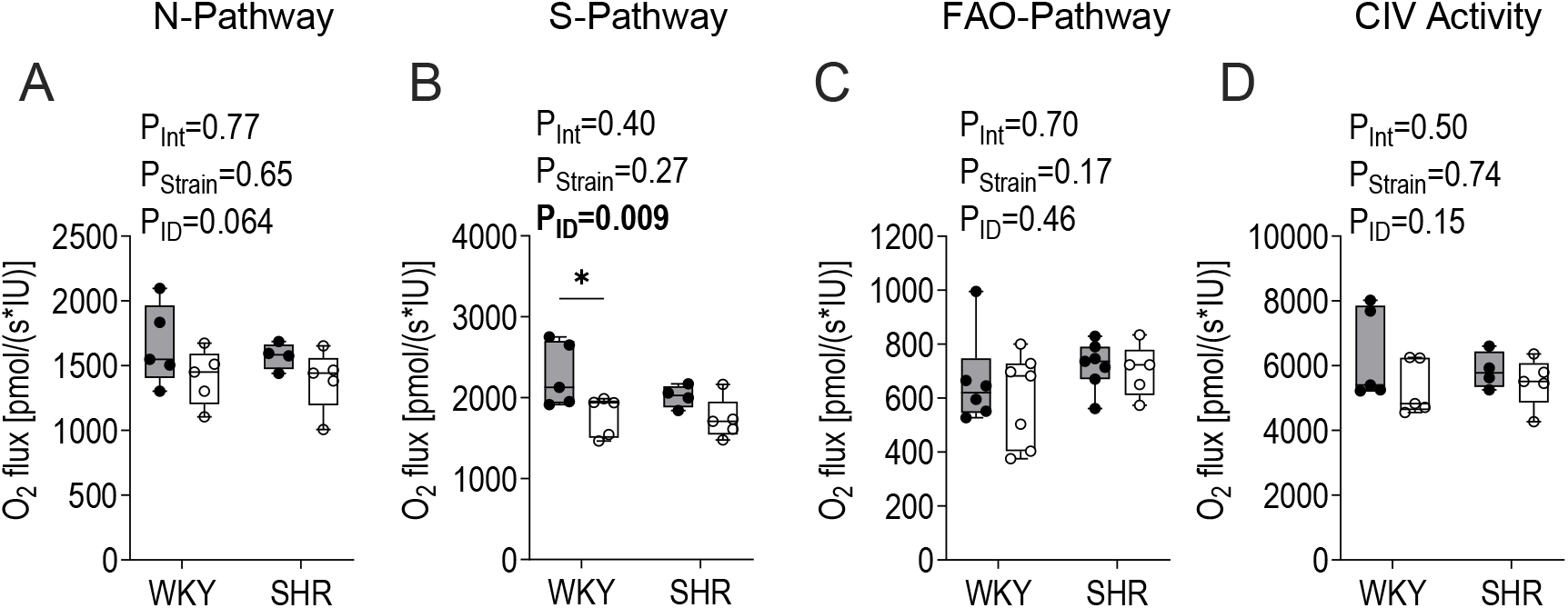
Iron deficiency (ID) differentially affects mitochondrial respiratory pathways in maternal cardiac tissue of Wistar Kyoto (WKY) and Spontaneously Hypertensive Rat (SHR) dams at gestational day 21. Oxygen flux normalized to citrate synthase activity (IU) is shown for (A) NADH-linked (N-Pathway) respiration, (B) succinate-linked (S-Pathway) respiration, (C) fatty acid oxidation-linked (FAO-Pathway), and (D) complex IV (CIV) activity. Data are shown as box-and-whisker plots depicting median (middle line), 25th and 75th percentiles (lower and upper edges) and minimum and maximum values (whiskers), with biological replicates overlaid (n=4-8/group for all panels). Comparisons were performed using 2-Way ANOVA followed by Holm-Šídák’s post hoc test; *P<0.05. Statistical outliers, identified using Grubb’s test, included one ID-SHR datapoint in panel C.

Renal cortical N- and S-pathway respiration were lower in SHR than WKY dams (**Fig. S2F, G)**. Cortical S-pathway respiration was further reduced by ID, with the effect most apparent in SHR dams (**Fig. S2G**). FAO-supported respiration and CIV activity were unaffected by strain or ID (**Fig. S2H, I**).

### Mitochondrial Dynamics

Maternal cardiac markers of fusion and fission were assessed to determine whether ID was associated with altered mitochondrial remodelling. The inner mitochondrial membrane (IMM) fusion protein optic atrophy 1 (OPA1) was quantified as its long (L-OPA1) and short (S-OPA1) isoforms, with the L-OPA1:S-OPA1 ratio used as an index of OPA1 processing (15). L-OPA1 abundance was unchanged across groups (**Fig. 3A, left**), whereas S-OPA1 levels were elevated in SHR relative to WKY dams (**Fig 3A, middle**), resulting in a lower L-OPA1:S-OPA1 ratio in SHR dams (**Fig. 3A, right**). Outer membrane fusion (OMM) proteins mitofusin 1 and 2 (MFN1 and MFN2) were also altered. ID reduced MFN1 abundance overall, with the greater reduction observed in ID-SHR relative to Ctl-SHR dams (**Fig. 3B**). MFN2 was lower in SHR compared to WKY dams and was further reduced in ID-SHR compared to Ctl-SHR dams (**Fig. 3C**). In contrast, the fission adaptor protein (FIS1) was unchanged across all groups (**Fig. 3D**). Phosphorylation of dynamin-related protein 1 (DRP1), expressed as the p-DRP1:total DRP1 ratio, was increased in ID dams, with the most pronounced effect in ID-WKY dams (**Fig. 3E**).

**Figure 3.**
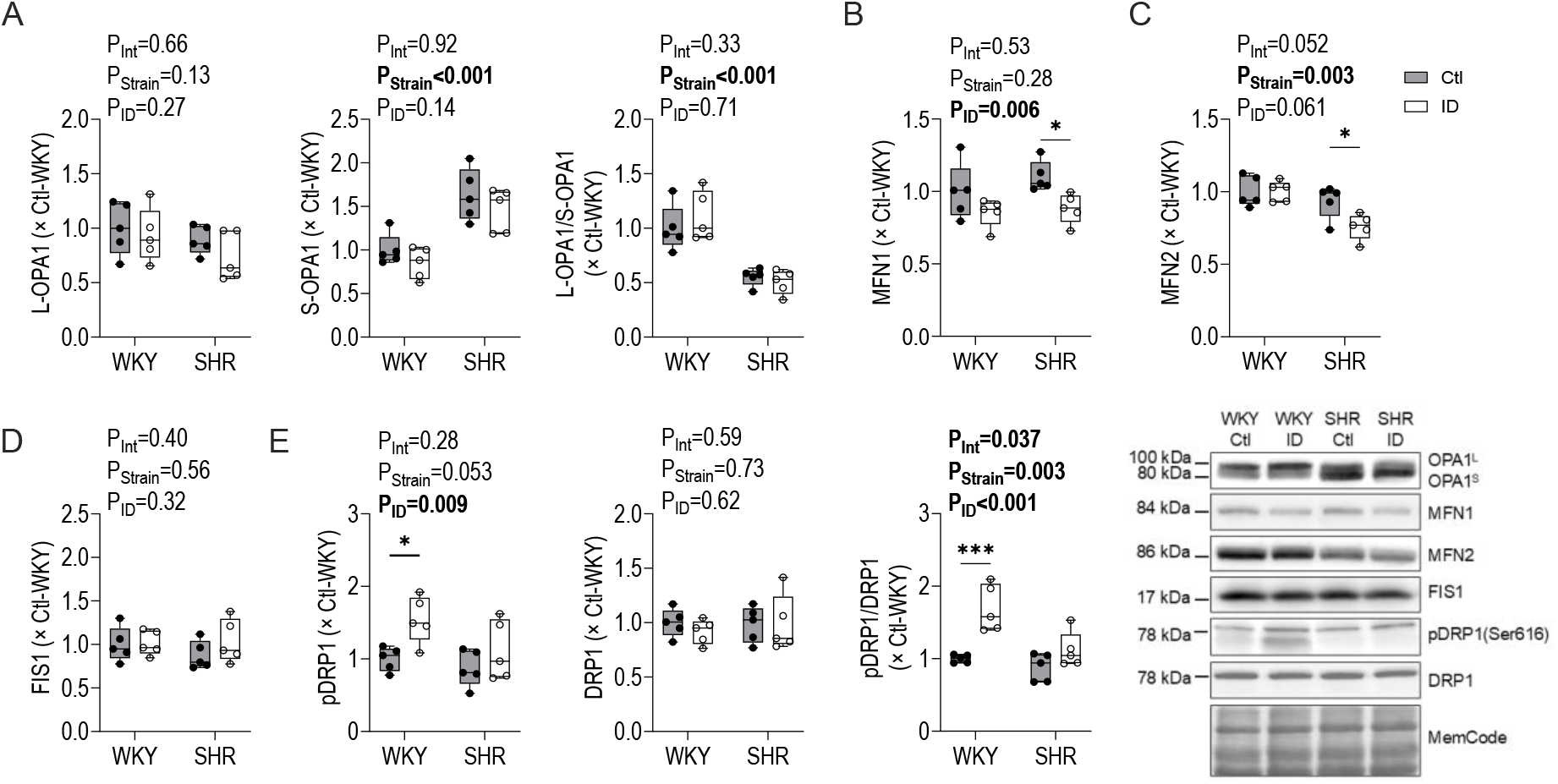
Iron deficiency (ID) induces strain-specific alterations in cardiac mitochondrial dynamics in Wistar Kyoto (WKY) and Spontaneously Hypertensive Rat (SHR) dams at gestational day 21. (A) Protein expression of long-form OPA1 (L-OPA1; left), short-form OPA1 (S-OPA1; middle), and the L-OPA1 to S-OPA1 ratio (right). (B) Mitofusin 1 (MFN1) protein expression. (C) Mitofusin 2 (MFN2) protein expression. (D) Fission 1 (FIS1) protein expression. (E) Phosphorylated DRP1 at Ser616 normalized to total DRP1 (pDRP1/DRP1). Representative immunoblots are shown on the right. Protein expression was normalized to MemCode total protein staining and expressed relative to the control (Ctl) WKY group. Data are shown as box-and-whisker plots depicting median (middle line), 25th and 75th percentiles (lower and upper edges) and minimum and maximum values (whiskers), with biological replicates overlaid (n=5/group in all panels). Comparisons were performed using 2-Way ANOVA followed by Holm-Šídák’s post hoc test; *P<0.05, ***P<0.001.

Given the observed alterations in mitochondrial ultrastructure and dynamics, markers of mitophagy were assessed to evaluate mitochondrial quality control pathways. The LC3-II:I ratio, a measure of autophagosome formation (16), was increased in ID dams, particularly in ID-SHR dams (**Fig. 4A**). BNIP3, a Parkin-independent mediator of mitophagy, was elevated in ID-SHR compared to Ctl-SHR, with no changes in ID-WKY dams (**Fig. 4B**). PINK1 expression, which was generally higher in SHR dams, was increased by ID in both strains (**Fig. 4C**). In contrast, Parkin and p62 expression were unchanged across all groups (**Fig. 4D, E**).

**Figure 4.**
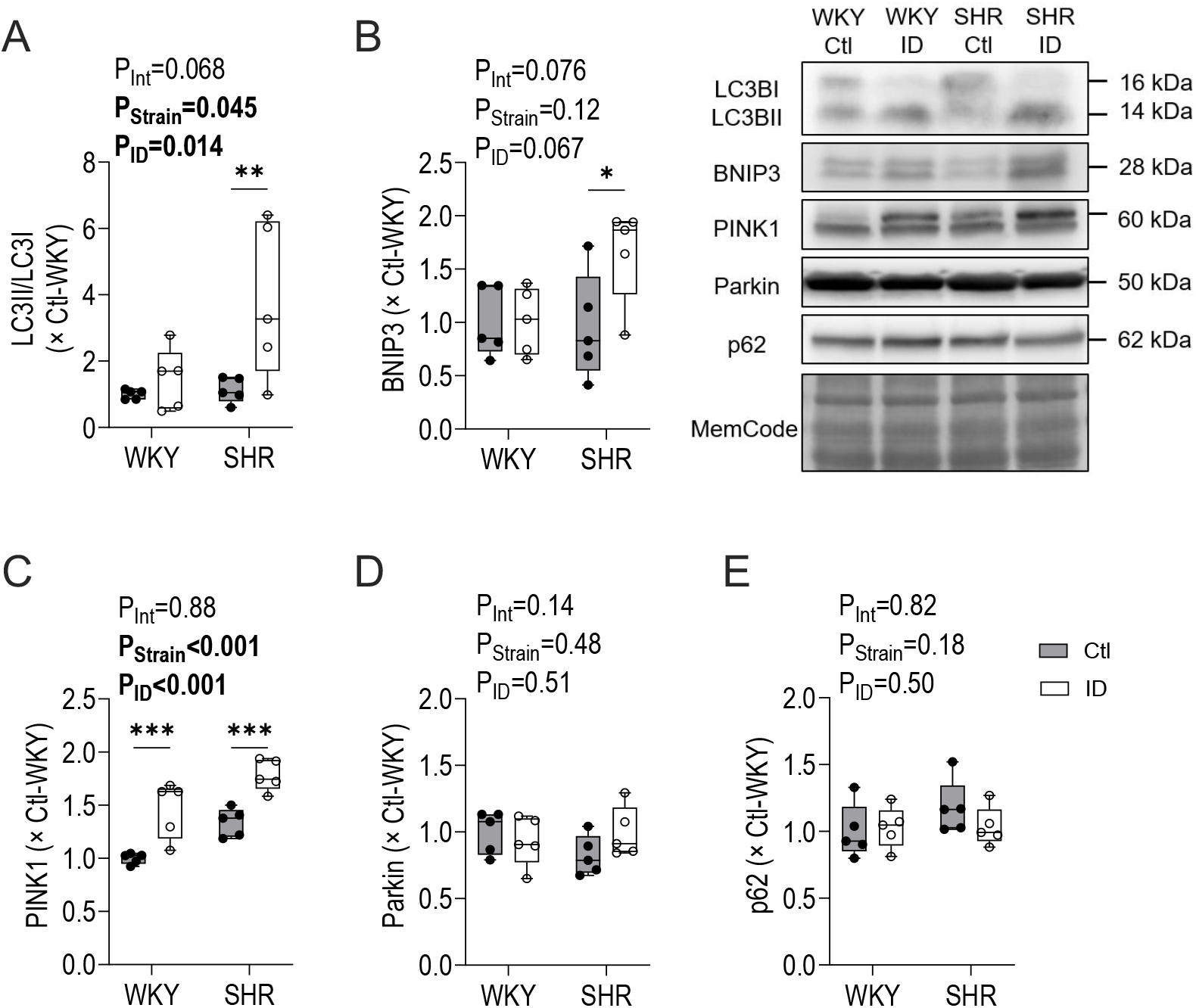
Iron deficiency (ID) induces strain-specific alterations in cardiac autophagy and mitophagy signalling in Wistar Kyoto (WKY) and Spontaneously Hypertensive Rat (SHR) dams at gestational day 21. (A) LC3II-to-LC3I ratio. (B) BNIP3 protein expression. (C) PINK1 protein expression. (D) Parkin protein expression. (E) p62 protein expression. Representative immunoblots are shown on the right. Protein expression was normalized to MemCode total protein staining and expressed relative to control (Ctl) WKY. Data are shown as box-and-whisker plots depicting median (middle line), 25th and 75th percentiles (lower and upper edges) and minimum and maximum values (whiskers), with biological replicates overlaid (n=5/group in all panels). Comparisons were performed using 2-Way ANOVA followed by Holm-Šídák’s post hoc test; *P<0.05, **P<0.01, ***P<0.001.

### Oxidative Stress

To assess redox regulation in the setting of mitochondrial remodelling, transcriptional markers of antioxidant defence and indices of oxidative stress were quantified (**Fig. 5**). Among the antioxidant genes assessed, catalase (*Cat*), cytosolic superoxide dismutase (*Sod*) 1, glutathione reductase (*Gsr*) and glutathione peroxidase 1 (*Gpx1*) all showed an interaction effect: ID increased expression in SHR hearts, whereas expression in WKY dams was either reduced or unchanged relative to their respective controls (**Fig. 5A**). In contrast, *Sod2* expression was unaffected by ID but remained higher overall in SHR compared with WKY dams (**Fig. 5A**). Malondialdehyde (MDA) expression, a marker of lipid peroxidation, showed a non-significant trending interaction (P_Int_=0.072; **Fig. 5B**), with elevations in ID-WKY relative to Ctl-WKY but no apparent effect of ID in SHR dams. Similarly, DHE fluorescence intensity exhibited an interaction effect: superoxide levels were elevated in ID-WKY compared with Ctl-WKY, whereas SHR dams exhibited elevated DHE signal irrespective of diet (**Fig. 5C**).

**Figure 5.**
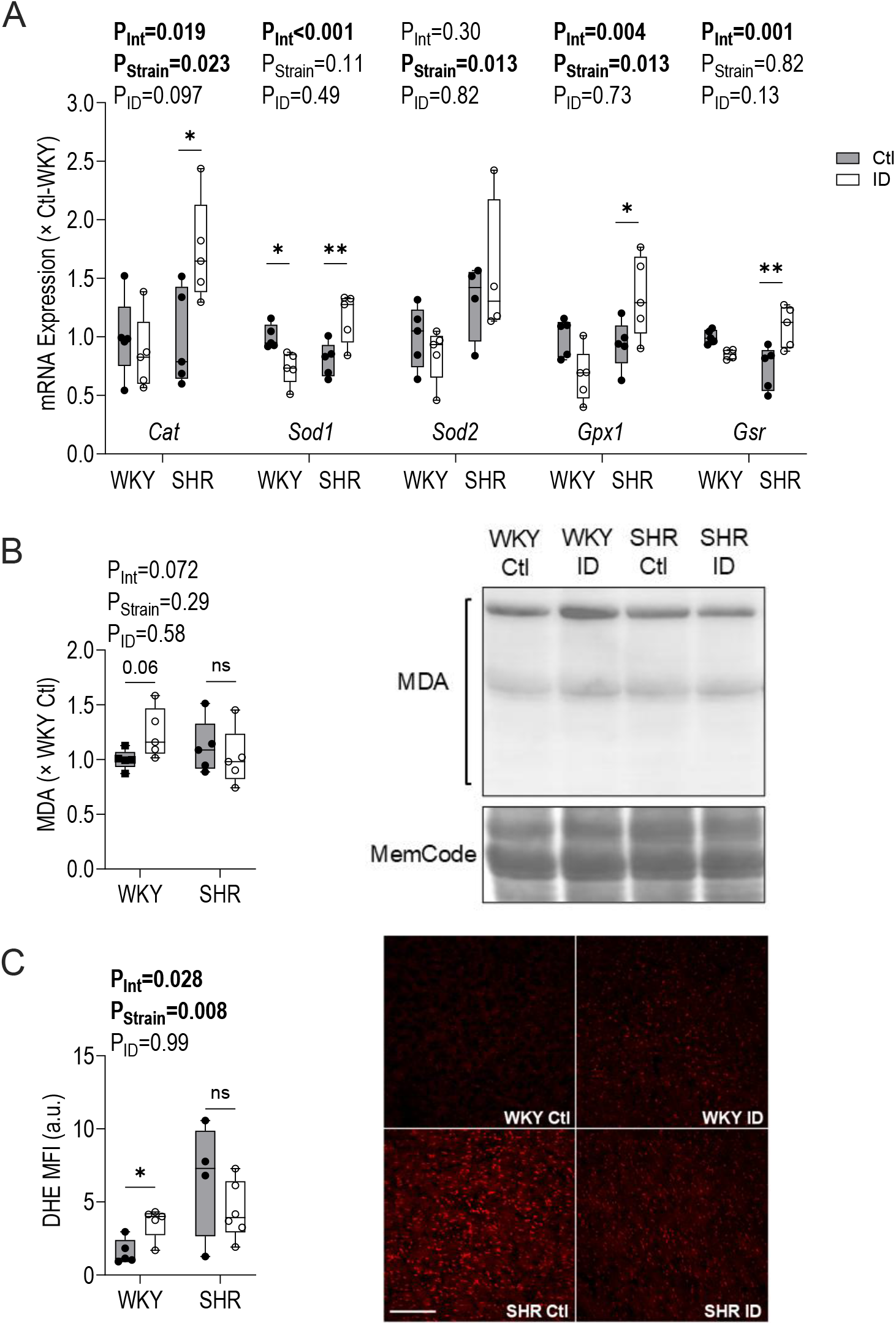
Iron deficiency (ID) induces strain-specific alterations in cardiac antioxidant enzyme expression while buffering oxidative stress in Wistar Kyoto (WKY) and Spontaneously Hypertensive Rat (SHR) dams at gestational day 21. (A) mRNA expression of catalase (*Cat*), superoxide dismutase 1 (*Sod1*), superoxide dismutase 2 (*Sod2*), glutathione peroxidase 1 (*Gpx1*), and glutathione reductase (*Gsr*), normalized to control (Ctl) WKY. (B) Malondialdehyde (MDA) protein expression was normalized to MemCode total protein staining and expressed relative to Ctl-WKY; representative immunoblots are shown to the right. (C) Dihydroethidium (DHE) mean fluorescence intensity (MFI), with representative images shown on the right (scale bar: 100 μm). Data are shown as box-and-whisker plots depicting median (middle line), 25th and 75th percentiles (lower and upper edges) and minimum and maximum values (whiskers), with biological replicates overlaid (n=4-7/group in all panels). Comparisons in panel A were performed using 2-Way ANOVA followed by Holm-Šídák’s post hoc test; *P<0.05, **P<0.01. Panels B and C were analyzed using non-parametric 2-way ANOVA, followed by within-strain comparisons using unpaired Student’s t-tests. Statistical outliers identified using Grubb’s test, included one Ctl-SHR and one ID-SHR datapoint in *Sod2* (panel A) and one ID-WKY datapoint in *Gsr* (panel A).

### Cellular Stress and Apoptosis

Cell survival signalling pathways were assessed to determine whether ID alters stress-responsive signalling in maternal cardiac tissue. The pAKT:AKT ratio showed an interaction effect (**Fig. 6A**), driven by increased phosphorylation in ID-WKY relative to Ctl-WKY dams, with no changes in SHR. The p-GSK3β:GSK3β ratio, a downstream effector of AKT, tended to be elevated in ID dams of both strains, although this did not reach statistical significance (P_ID_=0.052; **Fig. 6B**). NRF2 protein abundance was increased in ID dams independent of strain (**Fig. 6C**).

**Figure 6.**
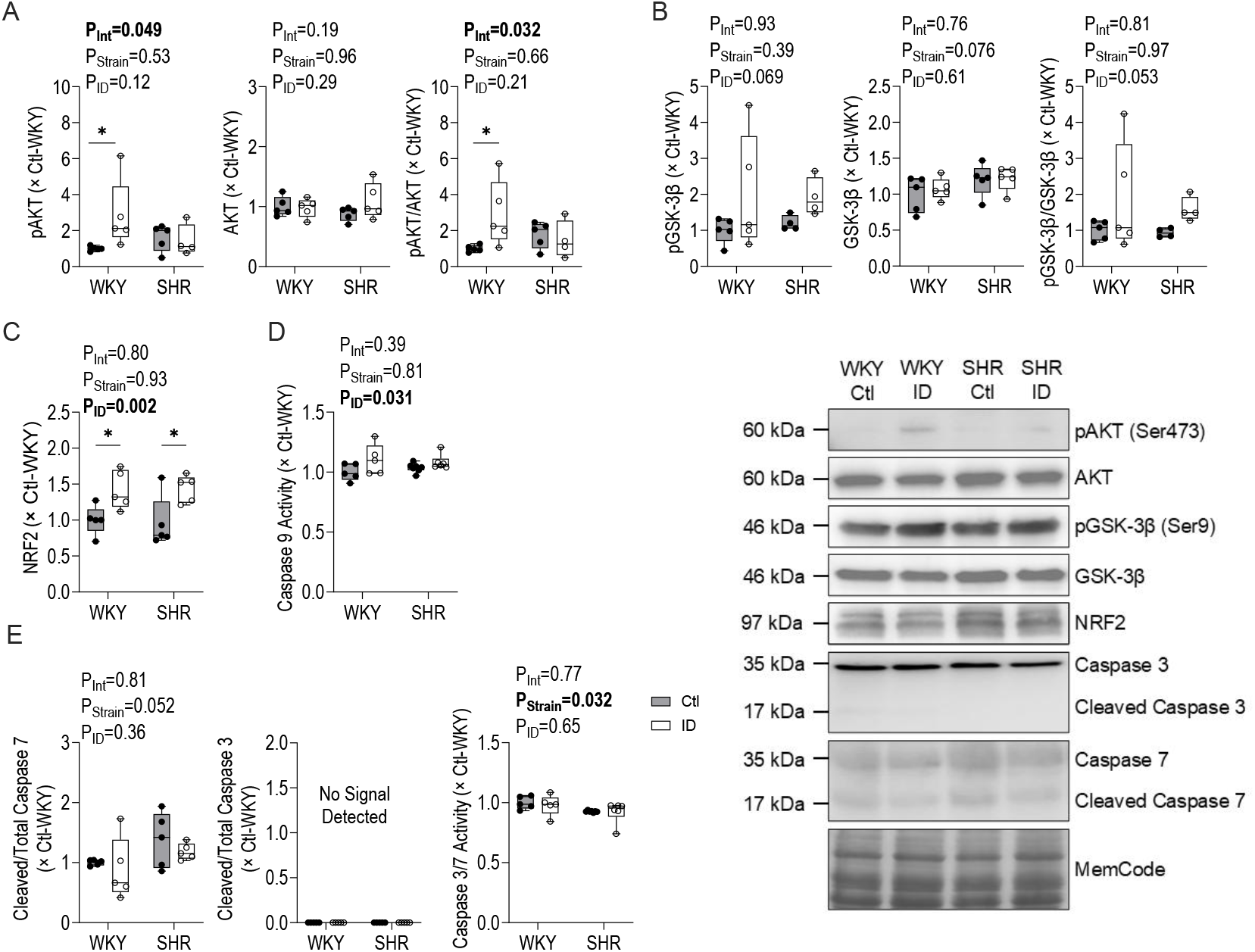
Iron deficiency (ID) engages AKT-dependent survival signalling without activation of mitochondrial-dependent apoptosis in the maternal heart of Wistar Kyoto (WKY) and Spontaneously Hypertensive Rat (SHR) dams at gestational day 21. (A) Protein expression of phosphorylated AKT at Ser473 (pAKT; left), total AKT (middle), and the pAKT/AKT ratio (right), normalized to the control (Ctl) WKY group. (B) Protein expression of phosphorylated GSK-3β at Ser9 (p GSK-3β; left), total GSK-3β (middle), and the pGSK-3β/GSK-3β ratio (right). (C) NRF2 protein expression. (D) Caspase 9 activity relative to Ctl-WKY. (E) Cleaved Caspase 7 normalized to total Caspase 7 protein expression (left), cleaved Caspase 3 normalized to total Caspase 3 protein expression (middle), and combined Caspase 3 and 7 activity (right) relative to the Ctl-WKY group. Protein expression was normalized to MemCode total protein staining and expressed relative to the Ctl-WKY group. Data are shown as box-and-whisker plots depicting median (middle line), 25th and 75th percentiles (lower and upper edges) and minimum and maximum values (whiskers), with biological replicates overlaid (n=4-7/group in all panels). Comparisons were performed using 2-Way ANOVA followed by Holm-Šídák’s post hoc test; *P<0.05. Statistical outliers, identified using Grubb’s test, included one ID-SHR datapoint in panel A and one Ctl-SHR and one ID-SHR datapoint in panel B.

Markers of apoptosis were subsequently examined to determine whether these signalling changes were accompanied by mitochondrial-dependent apoptotic pathways. Initiator caspase 9 activity was elevated in ID dams compared with controls (**Fig. 6D**). In contrast, the cleaved/total caspase 7 ratio was unaffected by ID or strain, although a non-significant trend towards a strain effect was observed (P=0.052; **Fig. 6E**). Cleaved caspase 3 was undetectable under the present experimental conditions, precluding calculation of cleaved/total caspase 3 ratio (**Fig. 6E**). Consistent with the absence of downstream apoptotic activation, executioner caspase 3/7 activity was lower in SHR than WKY dams, with no effect of ID in either strain (**Fig. 6E**).

## Discussion

We previously demonstrated that IDA lowered blood pressure and improved cardiac efficiency and ventricular-arterial coupling in pregnant rats, with these effects more pronounced in hypertensive dams (7). However, because iron is essential for mitochondrial energy metabolism, redox balance, and cellular stress responses, improved cardiovascular performance does not necessarily reflect preservation of myocardial bioenergetic health. In the present study, maternal ID produced a distinct mitochondrial phenotype in the pregnant heart: ultrastructural remodelling and constrained iron-dependent respiration were common in both strains, whereas the downstream engagement of mitochondrial dynamics, mitophagy, and redox signalling differed markedly between normotensive and hypertensive dams, The hypertensive heart showed reduced fusion signalling, elevated antioxidant transcriptional responses, and increased BNIP3-linked mitophagy, whereas the normotensive heart showed a fission-dominant remodelling pattern accompanied by oxidative stress. Together, these findings suggest that maternal ID does not simply overlay a uniform mitochondrial insult on the heart during pregnancy; rather, the mitochondrial response to iron restriction is shaped by the underlying cardiovascular state, and can coexist with the favourable haemodynamic profile previously observed in hypertensive pregnancy.

TEM analysis revealed that cardiac mitochondria from ID dams were larger and exhibited reduced cristae density. These ultrastructural alterations are consistent with our previous findings in neonatal ID rats (13) and with reports of mitochondrial swelling and cristae disorganization in iron deprivation and hypertension (12, 13, 17, 18) Although CS activity was unchanged across groups, the TEM findings indicate that the mitochondrial phenotype was nonetheless altered. While CS activity is commonly used as a proxy of mitochondrial content (19), enlarged mitochondria with reduced cristae density may maintain similar citric acid cycle enzymatic content while exhibiting altered inner membrane architecture. Consistent with altered mitochondrial regulation, expression of the mitochondrial biogenesis- and metabolism-associated genes *Ppargc1a* and *Ppara* was reduced in ID hearts. Thus, the present findings are more consistent with mitochondrial structural remodelling and altered regulation than with a simple loss of mitochondrial abundance. Similar mitochondrial swelling and cristae disorganization have also been reported in patients with heart failure and myocardial ID (20), supporting the notion that iron limitation may sensitize cardiac mitochondria to structural remodelling under cardiovascular stress (11). Importantly, renal cortical mitochondria showed no ID-associated changes in area or cristae density, while CS activity was similarly preserved. The preserved renal cortical ultrastructure despite comparable systemic iron restriction indicates that the pronounced mitochondrial remodelling observed in the heart is not a generalized consequence of maternal ID.

The structural phenotype was accompanied by selective changes in respiratory capacity. Iron is essential for mitochondrial respiration through its incorporation into iron-sulphur clusters and heme prosthetic groups of ETS complexes (10). Consistent with this role, succinate-pathway flux (through complex II) was reduced in ID hearts, with a tendency toward lower NADH-pathway respiration (through complex I) when normalized to CS activity, whereas downstream complex IV activity remained unaffected. Complexes I and II contain multiple Fe–S centres and may be especially sensitive to reduced iron availability, often showing early reductions in activity during iron deprivation (13, 21). Prior iron deprivation studies have similarly demonstrated remodelling of the mitochondrial proteome, preferentially affecting iron-containing complexes while sparing downstream electron transport (11, 13, 22). Additionally, FCRs were unchanged across groups, indicating that the relative contribution of each pathway to total respiratory flux was maintained despite lower complex II-linked respiratory capacity. These pathway specific changes are consistent with functional limitations at iron-dependent sites within the ETS. Notably, S-pathway respiration was also reduced in renal cortical tissue from ID-SHR dams, identifying complex II as a shared point of iron vulnerability across both tissues. Taken together with the absence of ultrastructural changes in the renal cortex, these observations suggest that the compounded metabolic demands of hypertensive pregnancy render the maternal myocardium particularly vulnerable to iron restriction, beyond the effects of ID alone on complex II activity that are apparent in other high-demand tissues.

Notably, fatty acid-supported respiration remained preserved in ID dams, indicating that maternal ID did not impair lipid-supported respiratory capacity. This finding is particularly relevant in late gestation, a metabolic state characterized by increased maternal insulin resistance and greater reliance on fatty acid utilization to support elevated energetic demand (23, 24). The preservation of FAO alongside reduced S-pathway respiration suggests that maternal ID was associated with pathway-specific respiratory constraints rather than a generalized suppression of oxidative metabolism. Maintenance of FAO capacity may also contribute to the preservation of maternal cardiac performance during IDA, by supporting ATP production despite modest impairments in iron-dependent ETS pathways. However, *Ppargc1a* and its metabolic partner *Ppara* were reduced in ID hearts, indicating that preserved fatty acid-supported respiration occurred despite reduced transcriptional support for mitochondrial biogenesis and metabolism (25). Although reduced PPARα expression is often associated with diminished FAO and increased reliance on glucose metabolism in hypertrophic hearts (25, 26), the preserved FAO-supported respiration observed here suggests that transcriptional changes may precede measurable functional impairments in lipid oxidative capacity.

Mitochondria are highly dynamic organelles that continuously undergo fusion and fission, processes regulating mitochondrial structure, resource exchange, and segregation of damaged regions (15, 27). Notably, alterations in mitochondrial dynamic proteins observed here were strain dependent. OPA1 processing differed primarily by strain, with SHR dams exhibiting increased S-OPA1 abundance compared to WKY dams (15). In contrast, ID was associated with reduced OMM fusion proteins, particularly in SHR dams, where ID-SHR hearts had lower MFN1 and MFN2 than Ctl-SHR hearts. Consistent with this, male SHR cardiomyocytes have previously been reported to exhibit reduced MFN2 and OPA1, consistent with reduced fusion reserve and limited mitochondrial adaptability (28). Given the established role of OPA1 in cristae maintenance (15, 27, 29), the altered OPA1 profile observed in SHR dams may have contributed to the more pronounced ultrastructural disorganization detected by TEM. However, S-OPA1 has also been implicated in mitochondrial protection during oxidative stress, limiting superoxide accumulation and Ca^2+^-induced mitochondrial permeability transition susceptibility (30). Thus, increased S-OPA1 may reflect a stress-adaptive response in SHR dams. Nevertheless, mitochondrial network integrity requires coordinated OPA1-dependent IMM fusion with OMM fusion proteins MFN1 and MFN2 (31, 32). The concurrent reductions in MFN1 and MFN2 in ID-SHR hearts therefore suggest that iron restriction constrained coordinated fusion-associated signalling, potentially contributing to the structural and bioenergetic remodelling observed.

Maternal ID also altered fission-associated signalling, reflected by increased DRP1 phosphorylation that was most pronounced in ID-WKY dams. In other models, iron deprivation preferentially activates DRP1-associated fission and promotes mitochondrial fragmentation without reducing fusion machinery, suggesting that fission may represent an early or dominant remodelling response to ID (33). The preserved OPA1 and MFN2 levels in ID-WKY hearts may indicate greater plasticity in the normotensive heart, although reduced MFN1 still suggests some constraint on fusion signalling. Importantly, mitochondria remained enlarged on TEM, indicating that the structural phenotype is unlikely to result solely from altered dynamics and may instead reflect ID-associated stress-induced swelling occurring alongside fission responses (17). Overall, these findings suggest that maternal IDA did not produce a uniform mitochondrial dynamics profile across strains. Rather, maternal IDA was associated with a greater fission response in WKY hearts and a more fusion constrained profile in SHR hearts.

The alterations in mitochondrial dynamics and reductions in iron-dependent respiration suggest increased mitochondrial stress, potentially increasing reliance on mitochondrial quality-control pathways. The increased LC3-II:LC3-I ratios and BNIP3 expression in ID-SHR compared to Ctl-SHR dams are consistent with engagement of BNIP3-linked mitophagy signalling (16). BNIP3 is closely associated with hypoxic and metabolic stress signalling and has been implicated in receptor-mediated mitochondrial clearance in cardiomyocytes (16). This response was most pronounced in ID-SHR dams, which also exhibited constrained fusion signalling, suggesting greater engagement of mitochondrial quality-control pathways in the iron-restricted hypertensive heart. Although PINK1 abundance was elevated in ID dams, the lack of change in Parkin and p62 expression suggests that this response reflects early mitochondrial stress sensing rather than PINK1-Parkin-dependent mitophagy, as PINK1 accumulates on damaged or depolarized mitochondria upstream of Parkin recruitment (34).

Despite these signs of mitochondrial stress, the downstream redox consequences of maternal IDA were strain specific. In normotensive WKY dams, ID was associated with reduced or unchanged expression of several antioxidant genes, increased DHE fluorescence and a trend towards higher MDA expression, collectively consistent with increased oxidative stress. ID has been shown to reduce expression of key antioxidant enzymes, including *Cat*, *Gpx1*, and *Sod1*, supporting antioxidant gene reductions observed in ID-WKY hearts (35). SHR dams exhibited higher overall DHE fluorescence, but this was not further increased by ID. Instead, ID-SHR hearts showed elevated antioxidant transcriptional responses, while *Sod2* expression remained higher in SHR irrespective of diet. One possible explanation is that the preexisting oxidative stress (i.e. that which was present before onset of IDA) had already induced an antioxidant response, thereby buffering the additional oxidative burden imposed by ID (36). Consistent with this, prior studies have shown that the SHR myocardium exhibits a higher basal ROS burden and elevated myocardial MnSOD activity relative to WKY (37, 38). As MnSOD/SOD2 is the principal mitochondrial superoxide scavenger (39), and cardiomyocyte-specific SOD2 deficiency increases superoxide and lipid peroxidation (40), the higher *Sod2* abundance in SHR hearts may have helped limit overt oxidative damage despite elevated ROS levels.

Despite evidence of mitochondrial and redox stress, maternal ID did not result in overt downstream apoptotic activation. Cleaved caspase-3 remained undetectable under the present experimental conditions, and the cleaved:total caspase-7 ratio was unchanged. Despite increased caspase 9 activity, downstream caspase 3/7 activity remained unaffected by ID. Notably, SHR dams showed a non-significant trend toward higher cleaved caspase 7; given the elevated inflammatory signalling reported in SHR, this may reflect strain-related stress signalling rather than definitive apoptotic execution (41, 42). AKT phosphorylation increased in ID-WKY dams, with a non-significant trend toward increased GSK-3β phosphorylation, consistent with engagement of pro-survival signalling that may restrain mitochondrial permeability transition and downstream caspase activation (43). Increased NRF2 abundance in both strains further supports activation of a broader cytoprotective response. In terminally differentiated cardiomyocytes, partial activation of apoptotic signalling does not invariably culminate in cell death and has been observed during stress adaptation and cardiomyocyte remodelling (44). Together, these findings suggest that ID was associated with mitochondrial stress and apoptotic susceptibility, while concurrent activation of pro-survival and cytoprotective pathways may have limited progression to downstream apoptotic execution. However, alternative cell death pathways such as pyroptosis or ferroptosis cannot be excluded (45).

Although our previous work demonstrated that maternal IDA improved myocardial efficiency and ventricular-arterial coupling in hypertensive pregnancy, the present findings suggest that these functional gains do not reflect enhanced mitochondrial bioenergetics (7). Cardiac efficiency, calculated as the ratio of stroke work to total pressure-volume area, is a load-dependent index that was improved in ID-SHR dams primarily through a reduction in potential energy alongside preserved stroke work (7). It was inferred that this redistribution might also reflect a proportional reduction in non-contractile myocardial energy expenditure. The present data support this interpretation: reduced PGC1α/PPARα expression with lower complex II-supported respiration and a trend toward reduced complex I-supported respiration indicate constrained metabolic capacity in the maternal heart. Together, these findings suggest that the apparent efficiency gain may reflect reduced metabolic demand rather than enhanced bioenergetic function, although contribution from reduced afterload cannot be excluded. Concurrent activation of stress-response and pro-survival pathways may also explain why these mitochondrial constraints did not progress to overt cardiac injury within the gestational window examined.

Several caveats warrant consideration for clinical translation. First, the trajectory of these changes may depend on the duration and the severity of anaemia, as well as the baseline hypertensive pathology. The relatively short gestational window examined here may capture an early compensatory phase, whereas the longer exposure typical of human pregnancy could lead to greater pathology. Importantly, these mitochondrial changes emerged despite only modest baseline hypertension and mild IDA, suggesting that maternal cardiac mitochondrial stress may arise before severe cardiovascular disease is apparent. Second, female SHRs exhibit greater antioxidant defences than males, likely driven by oestrogen signalling and associated with greater cardioprotection (46, 47). The relatively young dams used here (15–16-week-old) may therefore retain compensatory pathways, whereas prolonged pre-gestational hypertension could promote pathological myocardial remodelling (48). Finally, although the *Tfrc* and *Hamp1* expression changes previously reported in this cohort (7), confirm functional ID as the underlying cause of anaemia, the present study cannot fully dissociate the mitochondrial effects of reduced iron availability from those of reduced oxygen delivery secondary to anaemia. The mitochondrial effects described here likely reflect interactions among iron-dependent processes, altered oxygen availability, and maternal metabolic adaptations, and disentangling these contributions will require additional testing in future studies.

In summary, these data identify maternal iron status as an important determinant of cardiac mitochondrial plasticity during hypertensive pregnancy and reveal that the mitochondrial response to maternal ID is not uniform across pregnancies with different underlying cardiovascular phenotypes. The normotensive heart engages fission-dominant remodelling and shows evidence of oxidative stress, whereas the hypertensive heart engages fusion suppression, mitophagy signalling, and antioxidant upregulation. Both patterns coexist with the favourable haemodynamic changes previously observed in maternal IDA, but neither reflects enhanced bioenergetic capacity. Whether these divergent mitochondrial phenotypes are reversible, sustainable, or ultimately maladaptive remains unknown and warrants further longitudinal investigation beyond gestation, particularly in relation to anaemia severity, iron repletion, and postpartum cardiovascular recovery. Defining how iron availability shapes myocardial mitochondrial function in HDP will be critical for understanding maternal cardiovascular risk and refining therapeutic approaches in high-risk pregnancies.

## Data Availability

All supporting data are included within the main article and the supplementary file.

## Acknowledgments

The authors acknowledge the University of Alberta Experimental Oncology Cell Imaging Facility for transmission electron microscopy imaging and the University of Alberta Cell Imaging Core for the use of their Leica TCS SP5 confocal microscope.

## References

1. Bisson, C., Dautel, S., Patel, E., Suresh, S., Dauer, P. and Rana, S. (2023) Preeclampsia pathophysiology and adverse outcomes during pregnancy and postpartum. Front Med (Lausanne). 10, 1144170. 10.3389/fmed.2023.1144170

2. Rosenberg, E.A. and Seely, E.W. (2024) Update on Preeclampsia and Hypertensive Disorders of Pregnancy. Endocrinol Metab Clin North Am. 53, 377–389. 10.1016/j.ecl.2024.05.012

3. 3. World Health Organization. Anaemia. [Accessed July 24 2026]. Available from: https://who.int/news-room/fact-sheets/detail/anaemia.

4. Stevens, G.A., Finucane, M.M., De-Regil, L.M., Paciorek, C.J., Flaxman, S.R., Branca, F. et al. (2013) Global, regional, and national trends in haemoglobin concentration and prevalence of total and severe anaemia in children and pregnant and non-pregnant women for 1995-2011: a systematic analysis of population-representative data. Lancet Glob Health. 1, e16–25. 10.1016/S2214-109X(13)70001-9

5. Benson, A.E., Shatzel, J.J., Ryan, K.S., Hedges, M.A., Martens, K., Aslan, J.E. et al. (2022) The incidence, complications, and treatment of iron deficiency in pregnancy. Eur J Haematol. 109, 633–642. 10.1111/ejh.13870

6. Woodman, A.G., Mah, R., Keddie, D.L., Noble, R.M.N., Holody, C.D., Panahi, S. et al. (2020) Perinatal iron deficiency and a high salt diet cause long-term kidney mitochondrial dysfunction and oxidative stress. Cardiovasc Res. 116, 183–192. 10.1093/cvr/cvz029

7. Rachid, J.R., Vu, J., Liu, S.N., Panahi, S., Badhan, N.S., Holody, C.D. et al. (2025) Effects of iron deficiency anaemia on maternal haemodynamics and cardiac function in pregnant spontaneously hypertensive rats. Cardiovasc Res. 121, 1956–1968. 10.1093/cvr/cvaf149

8. Olivetti, G., Quaini, F., Lagrasta, C., Ricci, R., Tosini, P., Capasso, J.M. et al. (1993) Effects of genetic hypertension and nutritional anaemia on ventricular remodelling and myocardial damage in rats. Cardiovasc Res. 27, 1316–1325. 10.1093/cvr/27.7.1316

9. Olivares, M., Araya, M., Pizarro, F. and Letelier, A. (2006) Erythrocyte CuZn superoxide dismutase activity is decreased in iron-deficiency anemia. Biol Trace Elem Res. 112, 213–220. 10.1385/BTER:112:3:213

10. Paul, B.T., Manz, D.H., Torti, F.M. and Torti, S.V. (2017) Mitochondria and Iron: current questions. Expert Rev Hematol. 10, 65–79. 10.1080/17474086.2016.1268047

11. Hoes, M.F., Grote Beverborg, N., Kijlstra, J.D., Kuipers, J., Swinkels, D.W., Giepmans, B.N.G. et al. (2018) Iron deficiency impairs contractility of human cardiomyocytes through decreased mitochondrial function. Eur J Heart Fail. 20, 910–919. 10.1002/ejhf.1154

12. Eirin, A., Lerman, A. and Lerman, L.O. (2014) Mitochondrial injury and dysfunction in hypertension-induced cardiac damage. Eur Heart J. 35, 3258–3266. 10.1093/eurheartj/ehu436

13. Holody, C.D., Woodman, A.G., Nie, C., Liu, S.N., Young, D., Wiedemeyer, A. et al. (2025) Perinatal iron deficiency alters the cardiac proteome and mitochondrial function in neonatal offspring. Am J Physiol Heart Circ Physiol. 328, H101–H112. 10.1152/ajpheart.00412.2024

14. Woodman, A.G., Mah, R., Keddie, D., Noble, R.M.N., Panahi, S., Gragasin, F.S. et al. (2018) Prenatal iron deficiency causes sex-dependent mitochondrial dysfunction and oxidative stress in fetal rat kidneys and liver. FASEB J. 32, 3254–3263. 10.1096/fj.201701080R

15. Tilokani, L., Nagashima, S., Paupe, V. and Prudent, J. (2018) Mitochondrial dynamics: overview of molecular mechanisms. Essays Biochem. 62, 341–360. 10.1042/EBC20170104

16. Shires, S.E. and Gustafsson, A.B. (2015) Mitophagy and heart failure. J Mol Med (Berl). 93, 253–262. 10.1007/s00109-015-1254-6

17. Dong, F., Zhang, X., Culver, B., Chew, H.G., Jr., Kelley, R.O. and Ren, J. (2005) Dietary iron deficiency induces ventricular dilation, mitochondrial ultrastructural aberrations and cytochrome c release: involvement of nitric oxide synthase and protein tyrosine nitration. Clin Sci (Lond). 109, 277–286. 10.1042/CS20040278

18. Hickey, A.J., Chai, C.C., Choong, S.Y., de Freitas Costa, S., Skea, G.L., Phillips, A.R., et al. (2009) Impaired ATP turnover and ADP supply depress cardiac mitochondrial respiration and elevate superoxide in nonfailing spontaneously hypertensive rat hearts. Am J Physiol Cell Physiol. 297, C766–774. 10.1152/ajpcell.00111.2009

19. Larsen, S., Nielsen, J., Hansen, C.N., Nielsen, L.B., Wibrand, F., Stride, N. et al. (2012) Biomarkers of mitochondrial content in skeletal muscle of healthy young human subjects. J Physiol. 590, 3349–3360. 10.1113/jphysiol.2012.230185

20. Zhang, H., Jamieson, K.L., Grenier, J., Nikhanj, A., Tang, Z., Wang, F. et al. (2022) Myocardial Iron Deficiency and Mitochondrial Dysfunction in Advanced Heart Failure in Humans. J Am Heart Assoc. 11, e022853. 10.1161/JAHA.121.022853

21. Haddad, S., Wang, Y., Galy, B., Korf-Klingebiel, M., Hirsch, V., Baru, A.M. et al. (2017) Iron-regulatory proteins secure iron availability in cardiomyocytes to prevent heart failure. Eur Heart J. 38, 362–372. 10.1093/eurheartj/ehw333

22. Rensvold, J.W., Ong, S.E., Jeevananthan, A., Carr, S.A., Mootha, V.K. and Pagliarini, D.J. (2013) Complementary RNA and protein profiling identifies iron as a key regulator of mitochondrial biogenesis. Cell Rep. 3, 237–245. 10.1016/j.celrep.2012.11.029

23. Liu, L.X. and Arany, Z. (2014) Maternal cardiac metabolism in pregnancy. Cardiovasc Res. 101, 545–553. 10.1093/cvr/cvu009

24. Schulman-Geltzer, E.B., Fulghum, K.L., Singhal, R.A., Hill, B.G. and Collins, H.E. (2024) Cardiac mitochondrial metabolism during pregnancy and the postpartum period. Am J Physiol Heart Circ Physiol. 326, H1324–H1335. 10.1152/ajpheart.00127.2024

25. Duncan, J.G. and Finck, B.N. (2008) The PPARalpha-PGC-1alpha Axis Controls Cardiac Energy Metabolism in Healthy and Diseased Myocardium. PPAR Res. 2008, 253817. 10.1155/2008/253817

26. Ismael, S., Purushothaman, S., Harikrishnan, V.S. and Nair, R.R. (2015) Ligand specific variation in cardiac response to stimulation of peroxisome proliferator-activated receptor-alpha in spontaneously hypertensive rat. Mol Cell Biochem. 406, 173–182. 10.1007/s11010-015-2435-x

27. Ong, S.B., Kalkhoran, S.B., Cabrera-Fuentes, H.A. and Hausenloy, D.J. (2015) Mitochondrial fusion and fission proteins as novel therapeutic targets for treating cardiovascular disease. Eur J Pharmacol. 763, 104–114. 10.1016/j.ejphar.2015.04.056

28. Quiroga, C., Mancilla, G., Oyarzun, I., Tapia, A., Caballero, M., Gabrielli, L.A. et al. (2020) Moderate Exercise in Spontaneously Hypertensive Rats Is Unable to Activate the Expression of Genes Linked to Mitochondrial Dynamics and Biogenesis in Cardiomyocytes. Front Endocrinol (Lausanne). 11, 546. 10.3389/fendo.2020.00546

29. Varanita, T., Soriano, M.E., Romanello, V., Zaglia, T., Quintana-Cabrera, R., Semenzato, M. et al. (2015) The OPA1-dependent mitochondrial cristae remodeling pathway controls atrophic, apoptotic, and ischemic tissue damage. Cell Metab. 21, 834–844. 10.1016/j.cmet.2015.05.007

30. Lee, H., Smith, S.B., Sheu, S.S. and Yoon, Y. (2020) The short variant of optic atrophy 1 (OPA1) improves cell survival under oxidative stress. J Biol Chem. 295, 6543–6560. 10.1074/jbc.RA119.010983

31. Chen, Y., Csordas, G., Jowdy, C., Schneider, T.G., Csordas, N., Wang, W. et al. (2012) Mitofusin 2-containing mitochondrial-reticular microdomains direct rapid cardiomyocyte bioenergetic responses via interorganelle Ca(2+) crosstalk. Circ Res. 111, 863–875. 10.1161/CIRCRESAHA.112.266585

32. 32. Cipolat, S., Martins de Brito, O., Dal Zilio, B. and Scorrano, L. (2004) OPA1 requires mitofusin 1 to promote mitochondrial fusion. Proc Natl Acad Sci U S A. 101, 15927–15932. 10.1073/pnas.0407043101

33. Zhong, X., Wu, Q., Wang, Z., Zhang, M., Zheng, S., Shi, F. et al. (2022) Iron deficiency exacerbates aortic medial degeneration by inducing excessive mitochondrial fission. Food Funct. 13, 7666–7683. 10.1039/d2fo01084d

34. Narendra, D.P., Jin, S.M., Tanaka, A., Suen, D.F., Gautier, C.A., Shen, J. et al. (2010) PINK1 is selectively stabilized on impaired mitochondria to activate Parkin. PLoS Biol. 8, e1000298. 10.1371/journal.pbio.1000298

35. Kurtoglu, E., Ugur, A., Baltaci, A.K. and Undar, L. (2003) Effect of iron supplementation on oxidative stress and antioxidant status in iron-deficiency anemia. Biol Trace Elem Res. 96, 117–123. 10.1385/BTER:96:1-3:117

36. Purdom-Dickinson, S.E., Lin, Y., Dedek, M., Morrissy, S., Johnson, J. and Chen, Q.M. (2007) Induction of antioxidant and detoxification response by oxidants in cardiomyocytes: evidence from gene expression profiling and activation of Nrf2 transcription factor. J Mol Cell Cardiol. 42, 159–176. 10.1016/j.yjmcc.2006.09.012

37. Csonka, C., Pataki, T., Kovacs, P., Muller, S.L., Schroeter, M.L., Tosaki, A. et al. (2000) Effects of oxidative stress on the expression of antioxidative defense enzymes in spontaneously hypertensive rat hearts. Free Radic Biol Med. 29, 612–619. 10.1016/s0891-5849(00)00365-8

38. Mi, C., Qin, X., Hou, Z. and Gao, F. (2019) Moderate-intensity exercise allows enhanced protection against oxidative stress-induced cardiac dysfunction in spontaneously hypertensive rats. Braz J Med Biol Res. 52, e8009. 10.1590/1414-431X20198009

39. Palma, F.R., He, C., Danes, J.M., Paviani, V., Coelho, D.R., Gantner, B.N. et al. (2020) Mitochondrial Superoxide Dismutase: What the Established, the Intriguing, and the Novel Reveal About a Key Cellular Redox Switch. Antioxid Redox Signal. 32, 701–714. 10.1089/ars.2019.7962

40. Sharma, S., Bhattarai, S., Ara, H., Sun, G., St Clair, D.K., Bhuiyan, M.S. et al. (2020) SOD2 deficiency in cardiomyocytes defines defective mitochondrial bioenergetics as a cause of lethal dilated cardiomyopathy. Redox Biol. 37, 101740. 10.1016/j.redox.2020.101740

41. Lamkanfi, M. and Kanneganti, T.D. (2010) Caspase-7: a protease involved in apoptosis and inflammation. Int J Biochem Cell Biol. 42, 21–24. 10.1016/j.biocel.2009.09.013

42. Miguel-Carrasco, J.L., Zambrano, S., Blanca, A.J., Mate, A. and Vazquez, C.M. (2010) Captopril reduces cardiac inflammatory markers in spontaneously hypertensive rats by inactivation of NF-kB. J Inflamm (Lond). 7, 21. 10.1186/1476-9255-7-21

43. Juhaszova, M., Zorov, D.B., Kim, S.H., Pepe, S., Fu, Q., Fishbein, K.W. et al. (2004) Glycogen synthase kinase-3beta mediates convergence of protection signaling to inhibit the mitochondrial permeability transition pore. J Clin Invest. 113, 1535–1549. 10.1172/JCI19906

44. Scheubel, R.J., Bartling, B., Simm, A., Silber, R.E., Drogaris, K., Darmer, D. et al. (2002) Apoptotic pathway activation from mitochondria and death receptors without caspase-3 cleavage in failing human myocardium: fragile balance of myocyte survival? J Am Coll Cardiol. 39, 481–488. 10.1016/s0735-1097(01)01769-7

45. Liu, D., Qin, H., Gao, Y., Sun, M. and Wang, M. (2024) Cardiovascular disease: Mitochondrial dynamics and mitophagy crosstalk mechanisms with novel programmed cell death and macrophage polarisation. Pharmacol Res. 206, 107258. 10.1016/j.phrs.2024.107258

46. Romero, M., Caniffi, C., Bouchet, G., Elesgaray, R., Laughlin, M.M., Tomat, A. et al. (2013) Sex differences in the beneficial cardiac effects of chronic treatment with atrial natriuretic Peptide in spontaneously hypertensive rats. PLoS One. 8, e71992. 10.1371/journal.pone.0071992

47. Pelzer, T., Jazbutyte, V., Hu, K., Segerer, S., Nahrendorf, M., Nordbeck, P. et al. (2005) The estrogen receptor-alpha agonist 16alpha-LE2 inhibits cardiac hypertrophy and improves hemodynamic function in estrogen-deficient spontaneously hypertensive rats. Cardiovasc Res. 67, 604–612. 10.1016/j.cardiores.2005.04.035

48. Bell, D., Kelso, E.J., Argent, C.C., Lee, G.R., Allen, A.R. and McDermott, B.J. (2004) Temporal characteristics of cardiomyocyte hypertrophy in the spontaneously hypertensive rat. Cardiovasc Pathol. 13, 71–78. 10.1016/S1054-8807(03)00135-2

