## Supplementary material for "Maternal iron deficiency elicits divergent cardiac mitochondrial responses in normotensive and hypertensive pregnancy"

**Supplementary Table S1.** Primer sequences for RT-qPCR.

| Gene [Gene ID] | | Primer Sequence (5'–3') |
|---|---|---|
| Beta-Actin [81822] | Fwd | CATGAAGATCAAGATCATTGCTCCT |
| | Rev | GCTGATCCACATCTGCTGGAA |
| Cat [24248] | Fwd | CGG ATT CCT GAG AGA GTG GTA CA |
| | Rev | TGT GGA GAA TCG GAC GGC AAT AG |
| Gpx1 [24404] | Fwd | GGC TCA CCC GCT CTT TAC C |
| | Rev | AAT GTC GTT GCG GCA CAC |
| Gsr [116686] | Fwd | GGG ATT GGC TGC GAT GAG AT |
| | Rev | TTC TGA AGA GGT AGG ATG AAT GGC G |
| Ppargc1a [83516] | Fwd | GGA GCA ATA AAG CAA AGA GCA |
| | Rev | GTG TGA GGA GGG TCA TCG TT |
| Ppara [25747] | Fwd | ATC ACC CGA GAG TTC CTA AA |
| | Rev | CCG ATC TCC ACA GCA AAT TAT |
| Sod1 [24786] | Fwd | GCA GAA GGC AAG CGG TGA |
| | Rev | GGT ACA GCC TTG TGT ATT GTC CC |
| Sod2 [24787] | Fwd | GTC TGT GGG AGT CCA AGG TT |
| | Rev | GTT CCT TGC AGT GGG TCC TGA TTA |

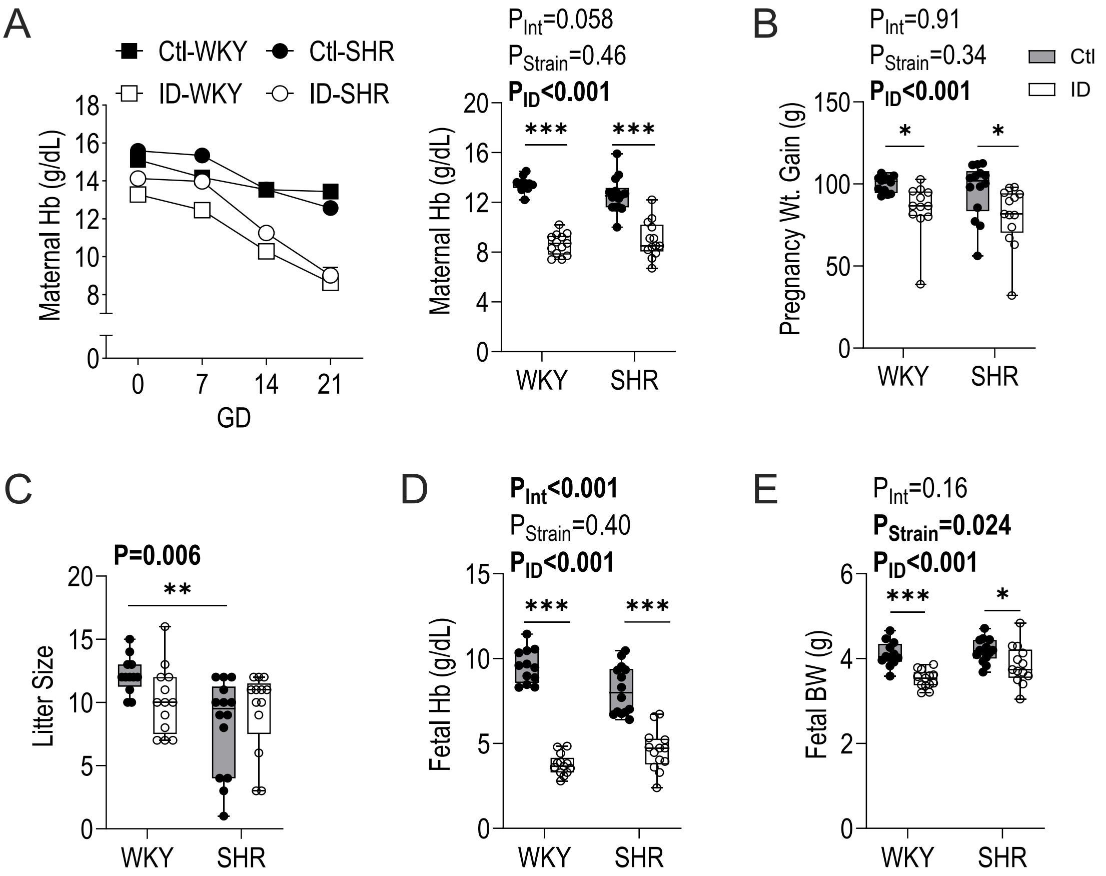


**Figure S1.** Maternal iron deficiency (ID) induces progressive anemia, reduces maternal weight gain, and differentially affects fetal hemoglobin, and fetal growth in Wistar Kyoto (WKY) and Spontaneously Hypertensive Rat (SHR) pregnancies at gestational day (GD) 21. (A) Maternal hemoglobin (Hb) levels throughout gestation (left panel) and at GD 21 (right panel). (B) Total pregnancy weight gain by GD21. (C) Litter size at GD21. (D) Fetal Hb concentration at GD21. (E) Fetal body weight (BW) at GD21. Data in panel A (left) are mean±SEM; remaining data are shown as box-and-whisker plots depicting median (middle line), 25th and 75th percentiles (lower and upper edges) and minimum and maximum values (whiskers), with biological replicates overlaid (n=12-14/group in all panels). Litter sizes were compared by Kruskal-Wallis test with Dunn's post hoc test. All other comparisons were made by 2-Way ANOVA; *$P<0.05$, **$P<0.01$, ***$P<0.001$ by Holm-Šídák's post hoc test.

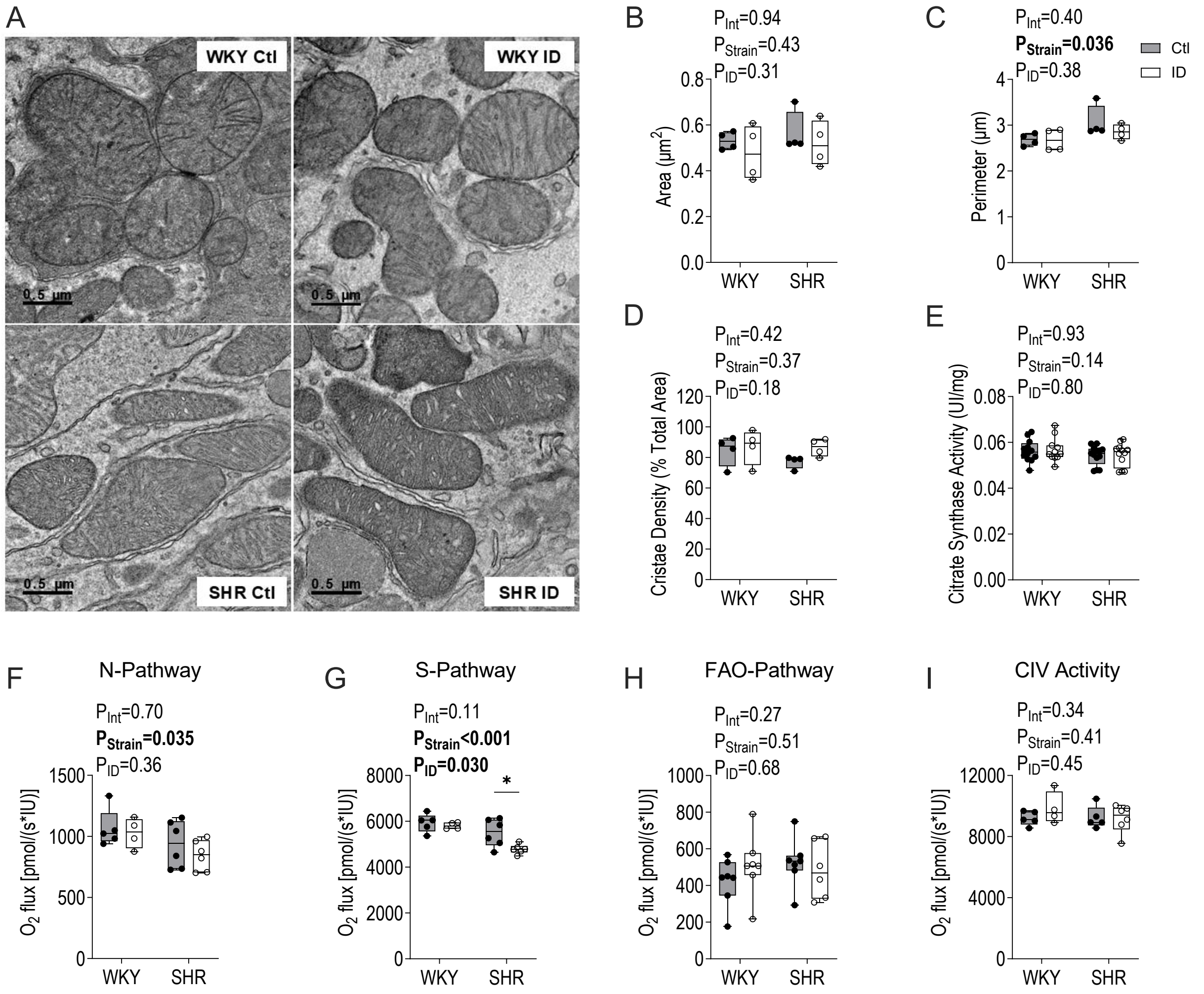


**Figure S2.** Kidney cortical mitochondrial ultrastructure and respiratory pathways in control (Ctl) and iron deficient (ID) Wistar Kyoto (WKY) and Spontaneously Hypertensive Rats (SHR) dams at gestational day 21. (A) Representative transmission electron microscopy images of renal cortex (scale bar: 0.5 µm). Quantification of (B) mitochondrial area, (C) mitochondrial perimeter, and (D) cristae density (% total area). (E) Citrate synthase activity (U/mg) in cortical tissue. High-resolution respirometry assessment of cortical mitochondrial oxygen flux through (F) NADH-linked (N-Pathway) respiration, (G) succinate-linked (S-Pathway) respiration, (H) fatty acid oxidation-linked (FAO-Pathway), and (I) complex IV (CIV) activity. Data are shown as box-and-whisker plots depicting median (middle line), 25th and 75th percentiles (lower and upper edges) and minimum and maximum values (whiskers), with biological replicates overlaid (n=4-13/group across panels). Comparisons were made by 2-Way ANOVA; *P<0.05 by Holm-Šídák's post hoc test.

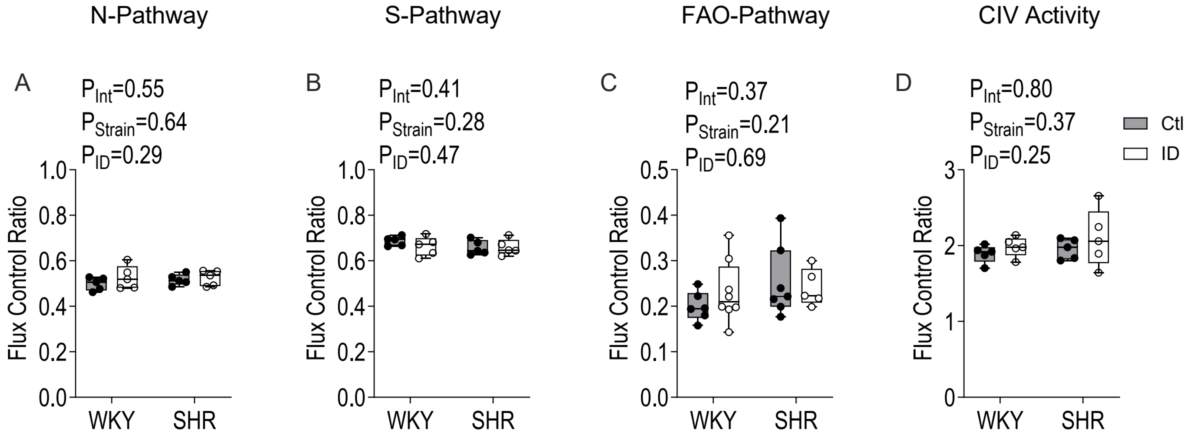


**Figure S3.** Iron deficiency (ID) minimally alters flux control ratios of mitochondrial respiratory pathways in Wistar Kyoto (WKY) and Spontaneously Hypertensive Rat (SHR) dams at gestational day 21. Flux control ratios calculated for (A) NADH-linked (N-Pathway) respiration, (B) succinate-linked (S-Pathway) respiration, (C) fatty acid oxidation (FAO), and (D) complex IV (CIV) activity. Data are shown as box-and-whisker plots depicting median (middle line), 25th and 75th percentiles (lower and upper edges) and minimum and maximum values (whiskers), with biological replicates overlaid (n=4-8/group in all panels). Comparisons were made by 2-Way ANOVA; *P<0.05 by Holm-Šídák's post hoc test.
